# pH-Dependent Evolution of Delafloxacin and Ciprofloxacin Resistance in *Pseudomonas aeruginosa* from cystic fibrosis (CF) and non-CF-patients

**DOI:** 10.64898/2026.09.08.750040

**Authors:** Niranjan Ireddy, Anja Bösch, Helena Seth-Smith, Philipp Kohler, Baharak Babouee Flury

## Abstract

Delafloxacin (DLX) is a novel fluoroquinolone with enhanced antibacterial activity in acidic environments, a property that may be advantageous for treating *Pseudomonas aeruginosa* infections in cystic fibrosis (CF), where airway surface liquid pH is typically reduced (pH 5.5–6.7). However, the propensity for resistance development and the underlying mechanisms in *P. aeruginosa* remain incompletely defined. We conducted serial passage experiments on six clinical *P. aeruginosa* isolates (three CF-and three non-CF-derived) exposed to sub-inhibitory concentrations of DLX or ciprofloxacin (CIP) at pH 6.0 and 7.3 over nine days. Susceptibility was assessed by broth microdilution (BMD), and resistance mechanisms were characterized by whole-genome sequencing (WGS), efflux pump expression analysis (qRT-PCR), and functional validation using CRISPR/Cas9-mediated genome editing and complementation assays. DLX minimal inhibitory concentrations (MICs) rose only 10.1-to 28.5-fold over 9 days, compared with 77.6-to 97.8-fold for CIP, indicating a substantially higher genetic barrier to resistance. This barrier was most pronounced under acidic conditions: only 38.9% of DLX-passaged samples crossed the resistance breakpoint, versus 94.4% at neutral pH, whereas CIP resistance reached 100% regardless of pH. Cross-resistance was asymmetric: exposure to DLX consistently selected for CIP cross-resistance (97.2% of samples), whereas exposure to CIP induced DLX cross-resistance efficiently at neutral pH but only partially under acidic conditions. A previously undescribed *gyrA* mutation (p.Ala51Val) conferred a 4-fold increase in DLX MIC when introduced by CRISPR/Cas9, and upregulation of the MexEF-OprN efflux pump, reversible by *mexS* complementation, emerged as a prominent resistance mechanism. Overall, DLX exhibited a markedly higher genetic barrier to resistance than CIP in *P. aeruginosa*, particularly under the acidic conditions characteristic of the CF airway. However, its use may co-select for CIP cross-resistance through efflux upregulation, underscoring the need for careful stewardship in CF.

## Introduction

*Pseudomonas aeruginosa* is an opportunistic pathogen that can cause severe infections, particularly in immunocompromised individuals and patients with cystic fibrosis (CF), in whom it commonly establishes persistent lung infection associated with substantial morbidity and mortality (1, 2). Its ability to adapt to diverse environments is driven by intrinsic and acquired antibiotic resistance mechanisms, as well as biofilm formation, which promote persistence and treatment failure (3–5).

Infections caused by *P. aeruginosa* can create acidic microenvironments (6). In biofilms, bacterial metabolic activity and the accumulation of acidic byproducts can further lower the local pH (7). Abscess fluid typically has a pH ranging from 5.5 to 7.2 (8).

Ciprofloxacin (CIP) is a first-line agent against *P. aeruginosa* and the only orally available option to date, but its activity is markedly reduced under acidic conditions, such as those present in abscesses and the airways of patients with cystic fibrosis (CF) (9–11).

Delafloxacin (DLX), currently approved for acute bacterial skin and skin structure infections (ABSSSI) and community-acquired pneumonia (CAP), is a dual-targeting, non-zwitterionic fluoroquinolone (12). Compared with CIP, its chemical properties promote enhanced bacterial uptake in both Gram-positive and Gram-negative pathogens and preserve activity under low-pH conditions such as those found in abscesses or infected tissues (13, 14).

DLX potency in acidic environments (12) may be particularly relevant for patients with CF, whose airways are often acidified (15). DLX has also shown superior in vitro activity against multidrug-resistant (MDR) *P. aeruginosa* compared with CIP in a CF sputum time-kill model (16).

However, data on DLX efficacy and resistance mechanisms in *P. aeruginosa* remain limited. In *Escherichia coli*, DLX resistance has been shown to develop more slowly than CIP resistance under acidic conditions and to be pH dependent, with mechanisms involving efflux pump upregulation and mutations in quinolone resistance determining region (QRDR) genes (17). Whether analogous or distinct mechanisms operate in *P. aeruginosa*, and whether the CF or non-CF origin of isolates influences resistance evolution, remains unknown. The present study therefore investigated the development of resistance and cross-resistance to DLX and CIP in *P. aeruginosa* at pH 6.0 and 7.3, characterized mechanisms of DLX resistance, and assessed its activity under acidic conditions.

## Results

### Resistance Evolution Kinetics

#### DLX demonstrates slower resistance evolution than CIP across CF and non-CF isolates

To compare the propensity for resistance development, we monitored MIC changes over 9 days of serial passage in sub-inhibitory antibiotic concentrations. Serial passage was restricted to 9 days to assess early resistance development under sustained sub-inhibitory selection pressure and to reflect the early clinical course of antibiotic exposure more closely, while minimizing prolonged laboratory adaptation and the accumulation of secondary compensatory changes. This time frame was sufficient to capture the initial divergence in resistance trajectories between DLX and CIP. At pH 6.0, CIP-exposed isolates showed a geometric mean MIC increase of 77.6-fold (95% CI: 27.0–223.1; *n* = 18), which was significantly greater than that observed in DLX-exposed isolates (10.1-fold; 95% CI: 5.1–19.9; *n* = 18; *p* = 0.0019, Wilcoxon signed-rank test). At pH 7.3, CIP resistance increased 97.8-fold (95% CI, 53.4–178.9), compared with 28.5-fold for DLX (95% CI, 16.7– 48.5; *p* = 0.011). No significant differences in resistance evolution were observed between CF and non-CF isolates for either antibiotic across both pH conditions (DLX, *p* = 0.99; CIP, *p* = 0.61, Mann-Whitney U test), indicating that the slower emergence of DLX resistance is independent of clinical origin (**Figure 1**).

**Figure 1:**
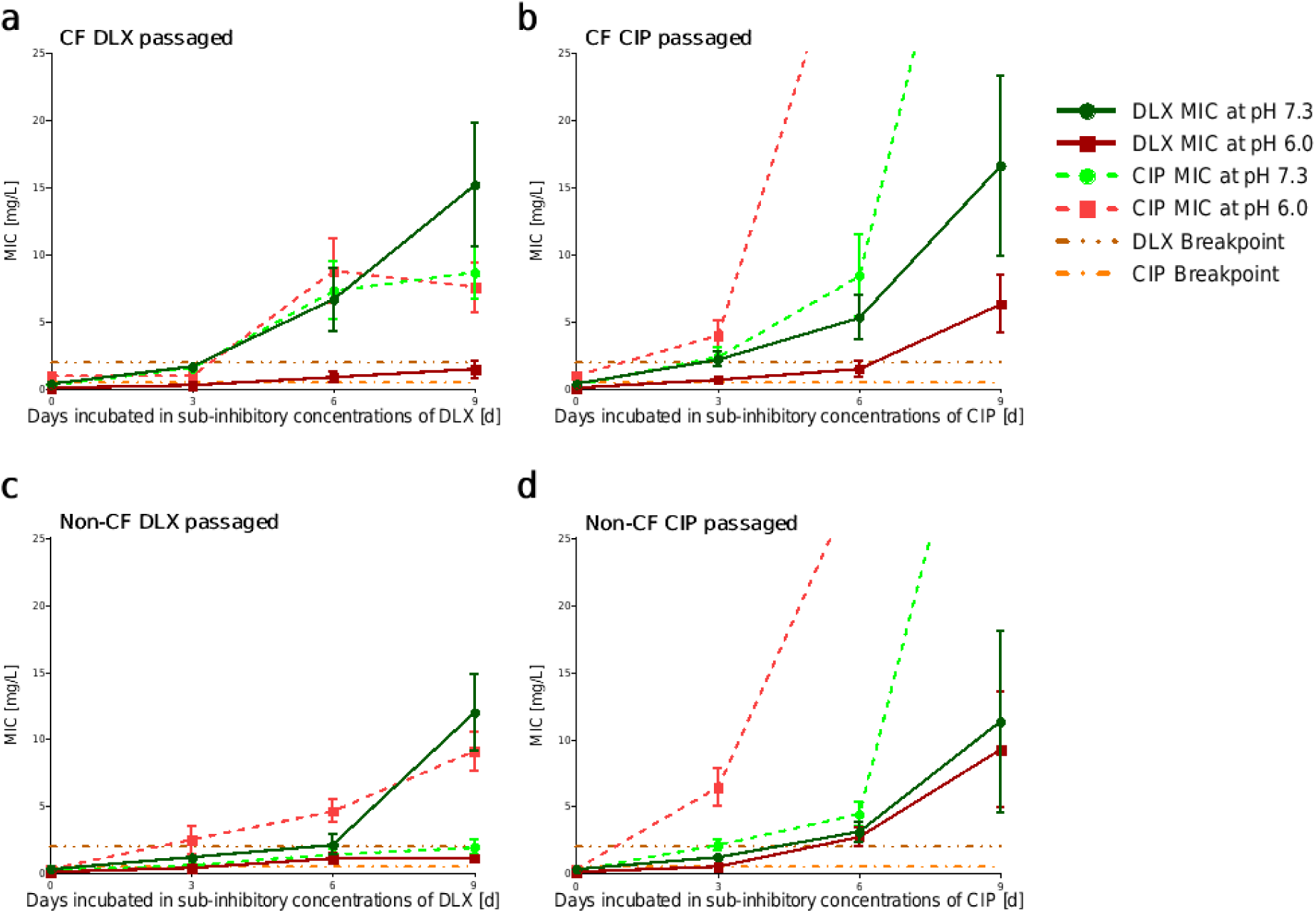
CF (a and b) and non-CF isolates (c and d) resistance evolution of *P. aeruginosa* passaged in subinhibitory (1/2 MIC) concentrations of delafloxacin (a and c) or ciprofloxacin (b and d) at pHs 7.3 and 6.0., horizontal dashed lines indicate clinically relevant breakpoints (EUCAST for CIP: >0.5 mg/L; FDA for DLX: ≥2 mg/L). Each trajectory represents the mean of one condition *n* = 18 biological replicates (3 clones × 6 parental isolates). (https://www.fda.gov/drugs/developmentresources/delafloxacin-injection-and-oral-products).

By day 9, all CIP-passaged isolates at pH 6.0 (18/18, 100%) exceeded the EUCAST resistance breakpoint (R > 0.5 mg/L), compared with 7/18 (38.9%) DLX-passaged isolates exceeding the FDA breakpoint (R ≥ 2 mg/L; McNemar’s exact *p* < 0.001). At pH 7.3, resistance was observed in 18/18 (100%) CIP and 17/18 (94.4%) DLX lineages (McNemar’s exact *p* = 1.0) indicating substantially slower resistance evolution under acidic conditions, whereas CIP resistance reached 100% at both pH levels.

As a sensitivity analysis to account for potential non-independence due to shared clonal ancestry, a population-averaged generalized estimating equation (GEE) model restricted to the DLX arm, clustered by parental strain and Mancl-DeRouen small sample corrected standard error adjusted for CF versus non-CF origin, confirmed the pH effect (odds ratio 27.4, 95% CI 2.80–267.5, *p* = 0.0044), while strain origin had no significant impact (*p* = 0.86)

At pH 6.0, CIP MICs increased 77.6-fold compared with 10.1-fold for DLX (*p* = 0.002), while at pH 7.3 the increases were 97.8-fold and 28.5-fold, respectively (*p* = 0.006). When susceptibility to the alternate agent was assessed, DLX-passaged isolates showed increases in CIP MIC of 11.3-fold (pH 6.0) and 14.3-fold (pH 7.3), whereas CIP-passaged isolates exhibited increases in DLX MIC of 48.9-fold (pH 6.0) and 21.0-fold (pH 7.3). Overall, the smaller fold-increase in MIC for DLX under selective pressure, compared with CIP at both pH levels, indicates a higher genetic barrier to resistance development (**Figure 2)**.

**Figure 2:**
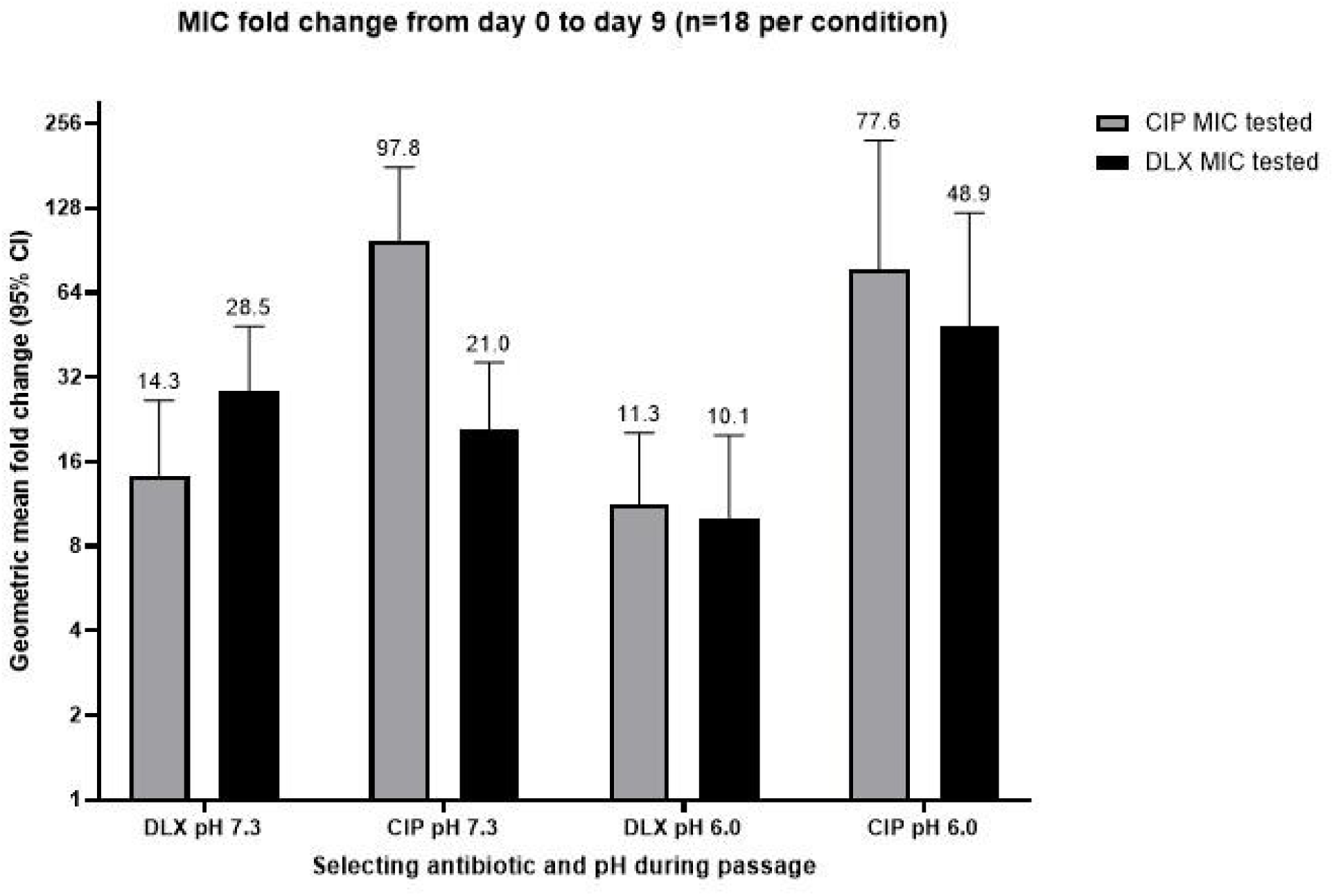
Quantitative comparison of resistance evolution to the selecting antibiotic (the drug an isolate was actually passaged under) versus cross-resistance to the non-selecting antibiotic (the drug it was never exposed to but was still tested against, to capture cross-resistance), across pH conditions. Grey bars show CIP MIC fold-change, black bars show DLX MIC fold-change.

### Cross-Resistance Patterns

#### DLX exposure more readily induces cross-resistance to CIP than vice versa, and weather this asymmetry is influenced by pH depends on the unit of independence

Among DLX-passaged isolates, 97.2% (35/36) developed CIP cross-resistance by day 9 (day 9 CIP MIC > 0.5 mg/L), with no significant difference between pH conditions: 100% (18/18) at pH 6.0 and 94.4% (17/18) at pH 7.3 (Fisher’s exact *p* = 1.0; McNemar’s exact *p* = 1.0; **Figure 1a, c**). This indicates that CIP cross-resistance emerged in nearly all DLX-passaged lineages at both pH levels.

In contrast, 86.1% (31/36) of CIP-passaged isolates developed DLX cross-resistance by day 9 (day 9 DLX MIC ≥ 2 mg/L), with a pH-dependent difference whose significance depends on the analysis: 72.2% (13/18) at pH 6.0 versus 100% (18/18) at pH 7.3 (Fisher’s exact *p* = 0.045; McNemar’s exact *p* = 0.0625).

Cross-resistance did not differ significantly when both selection directions were pooled across pH (DLX to CIP 97.2% vs CIP to DLX 86.1%; Fisher’s exact *p* = 0.199; McNemar’s exact *p* = 0.219). Accordingly, directional asymmetry in cross-resistance was more pronounced in acidic conditions, where CIP selection produced incomplete DLX cross-resistance, while DLX selection yielded CIP cross-resistance in nearly all lineages across both pH levels, this pattern is suggestive rather than established as the two analyses straddle at 0.05 threshold.

Whole-genome sequencing identified mutations in mexS, the repressor of the mexEF-oprN operon, in five derivatives four CIP-passaged and one DLX-passaged (Table 1). We therefore examined expression of MexEF-OprN and related efflux systems.

**Table 1:**
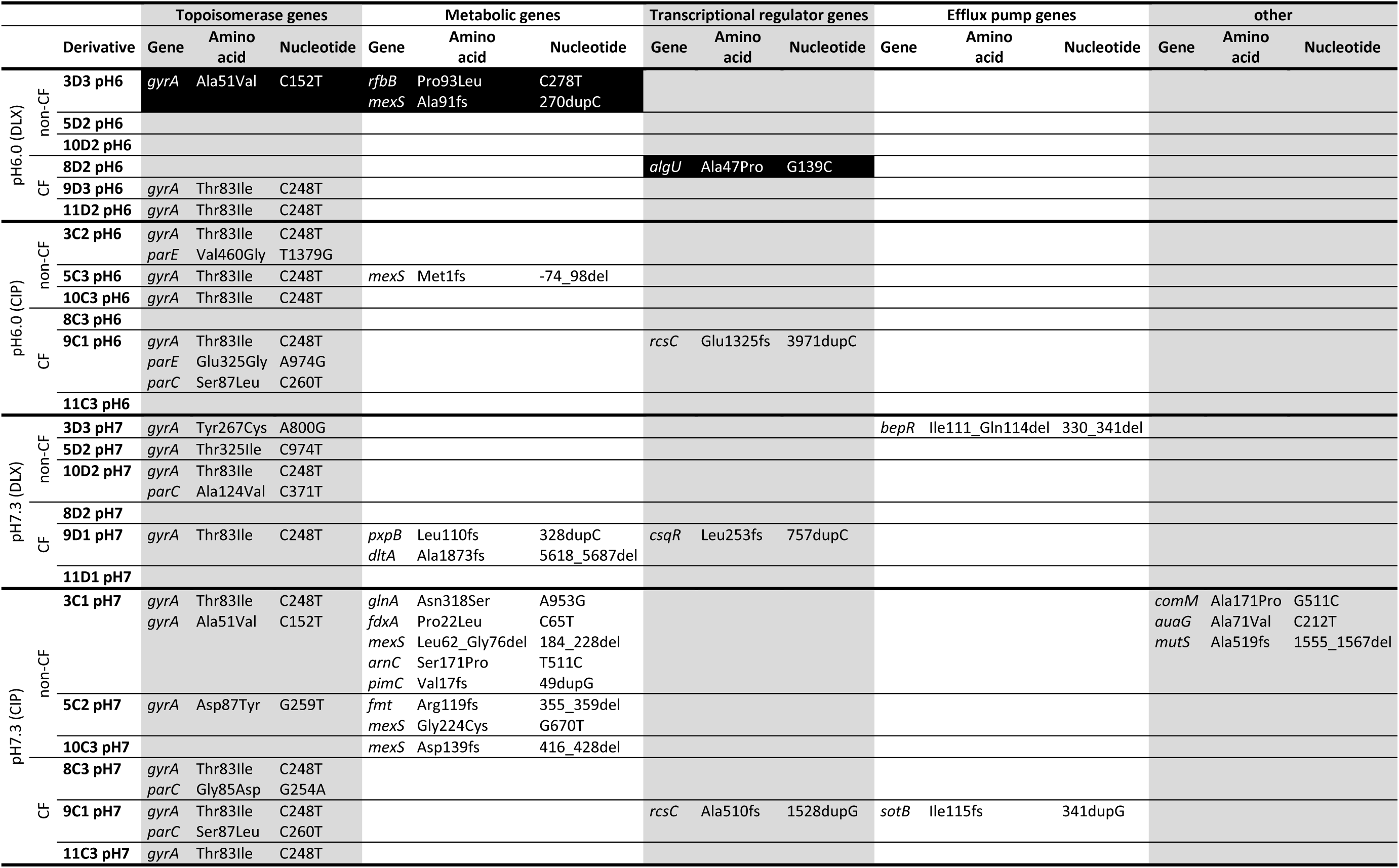
Non-synonymous SNPs identified by whole-genome sequencing in day 9 resistant derivatives relative to their day 0 parental strain (24 representative lineages), grouped by selection condition (antibiotic, pH) and clinical origin (CF/non-CF), and organized by gene functional category. DLX, delafloxacin; CIP, ciprofloxacin; CF, cystic fibrosis

### Molecular Mechanisms: Efflux Pump Expression

#### MexEF-OprN is the dominant efflux system upregulated under fluoroquinolone pressure, modulated by pH and antibiotic

Quantitative RT-PCR of 14 efflux-related genes in 24 resistant derivatives (**Supplemental Table 1 and 2**) showed strong upregulation of the MexEF-OprN structural and regulatory genes under selected conditions. In CIP-passaged isolates at pH 7.3, expression was much higher in non-CF than CF strains for *mexE*, *mexF,* and *oprN* (*mexE* 102.6 ± 81.6-fold vs 2.2 ± 1.3-fold, *mexF* 168.5 ± 149.9-fold vs 2.3 ± 1.6-fold, and *oprN* 27.4 ± 12.5-fold vs 2.1 ± 2.3-fold), and the same overall pattern was seen at pH 6.0, although the difference was smaller (*mexE* 74.2 ± 52.6-fold vs 6.8 ± 6.2-fold, *mexF* 143.6 ± 129.9-fold vs 9.5 ± 10.6-fold, and *oprN* 22.2 ± 20.8-fold vs 4.1 ± 4.3-fold), see **Figure 3**. Because only three CF and three non-CF isolates were available for each condition, none of the CF versus non-CF comparisons reached statistical significance; nevertheless, the direction and magnitude of change were consistent across all three genes, supporting a clear numerical trend toward stronger MexEF-OprN induction in non-CF isolates.

**Figure 3:**
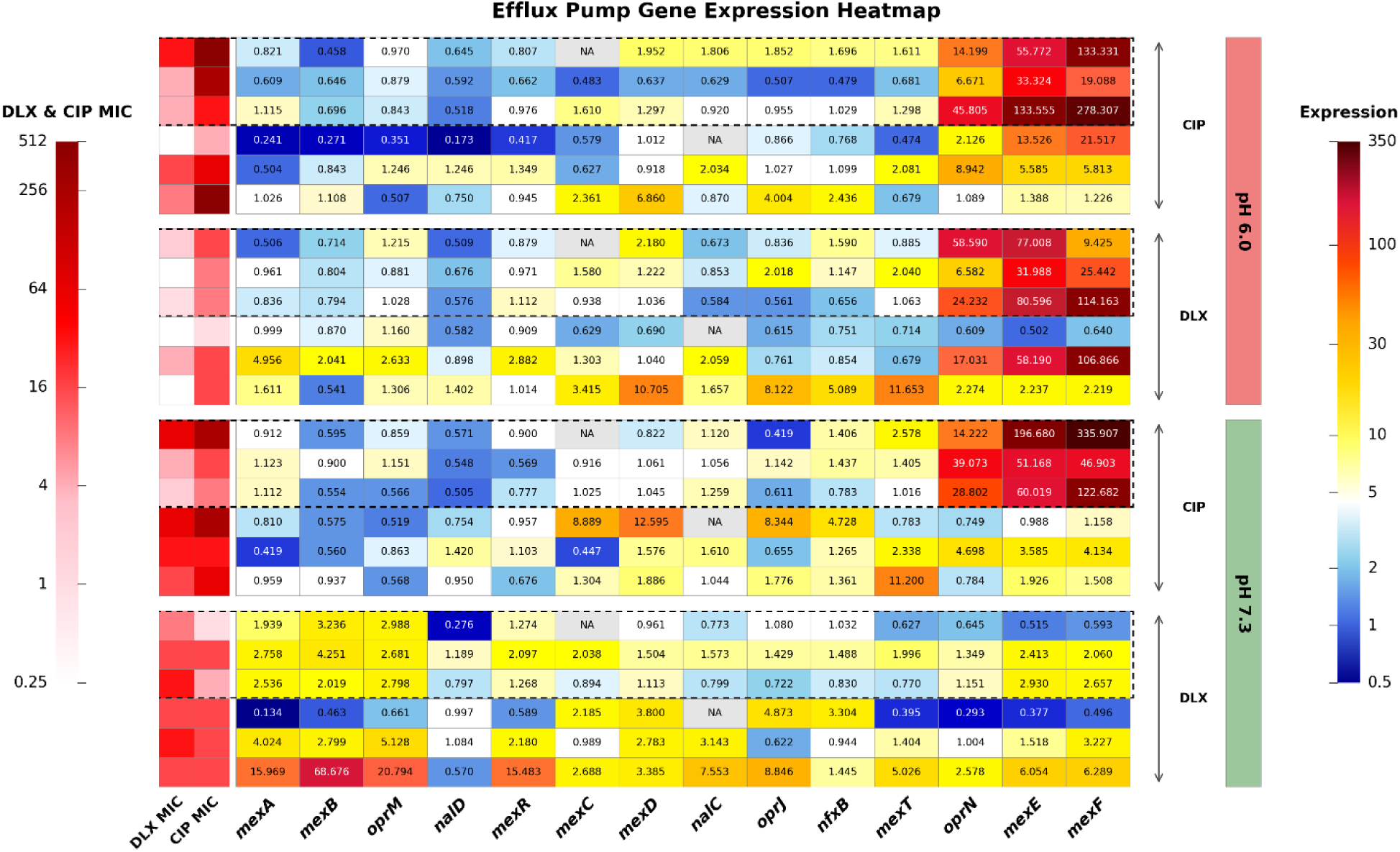
Expression heatmap of efflux pump genes and antibiotic susceptibility profiles in P. aeruginosa clinical isolates. Gene expression levels (fold-change relative to parental isolates) are shown with red indicating upregulation and blue indicating downregulation. Dotted box indicates non-CF isolates. DLX MIC and CIP MIC indicate delafloxacin and ciprofloxacin minimum inhibitory concentrations (μg/mL), respectively.

In contrast, DLX-passaged isolates showed a pattern more consistent with pH-dependent than origin-dependent upregulation across the same three genes. When pooled across origin, expression was higher at pH 6.0 than at pH 7.3 for all three genes (*mexE* 41.8 ± 35.7 vs 2.3 ± 2.1-fold, *p* = 0.13; *mexF* 43.1 ± 53.0 vs 2.6 ± 2.1-fold, *p* = 0.093; *oprN* 18.2 ± 21.8 vs 1.2 ± 0.8-fold, *p* = 0.065), indicating consistent differences that did not reach statistical significance at this sample size. By contrast, no significant CF/non-CF differences were observed within the DLX arm (*mexE p* = 0.18; *mexF p* = 0.59; *oprN p* = 0.31). Overall, the pH-associated effects were stronger than the origin-associated effects, with *mexF* and *oprN* showing the clearest trends.

MexAB-OprM was mostly near baseline across the 24 derivatives, with median fold-changes of 0.98 for *mexA*, 0.80 for *mexB*, and 1.00 for *oprM*. However, a small number of isolates showed marked overexpression, including one CF isolate at DLX pH 7.3 that accounted for the highest values observed. MexCD-OprJ showed the same general pattern: median expression remained close to baseline, but several isolates showed >2-fold increases, with *mexD* reaching 12.6-fold. Overall, these efflux systems were not uniformly induced across conditions, but instead showed sporadic, isolate-specific upregulation. Consistent with this, efflux gene expression was not strongly correlated with endpoint MIC, indicating that qPCR signal was better at identifying which genes were induced under specific conditions than at predicting resistance level.

### CRISPR/Cas9 Genome Engineering and Functional Validation

CRISPR/Cas9 genome engineering in the *P. aeruginosa* mPAO1 background targeted four candidate genes identified by SNP analysis as potential determinants of DLX resistance under acidic conditions: *gyrA*, *rfbB*, *mexS*, and *algU* (**Table 1**).

The *gyrA* C152T (p.Ala51Val) crispant showed a fourfold increase in DLX MIC relative to mPAO1 (**Figure 4**), with a corresponding twofold increase in CIP MIC, supporting this substitution as a moderate resistance determinant with a stronger effect on DLX (**Table 2a)**. In contrast, the *rfbB* C278T (p.Pro93Leu) crispant was associated with reduced MICs for both drugs under most conditions, consistent with a compensatory rather than a primary resistance role; the only exception was CIP at pH 6.0, where a twofold increase was observed.

**Figure 4:**
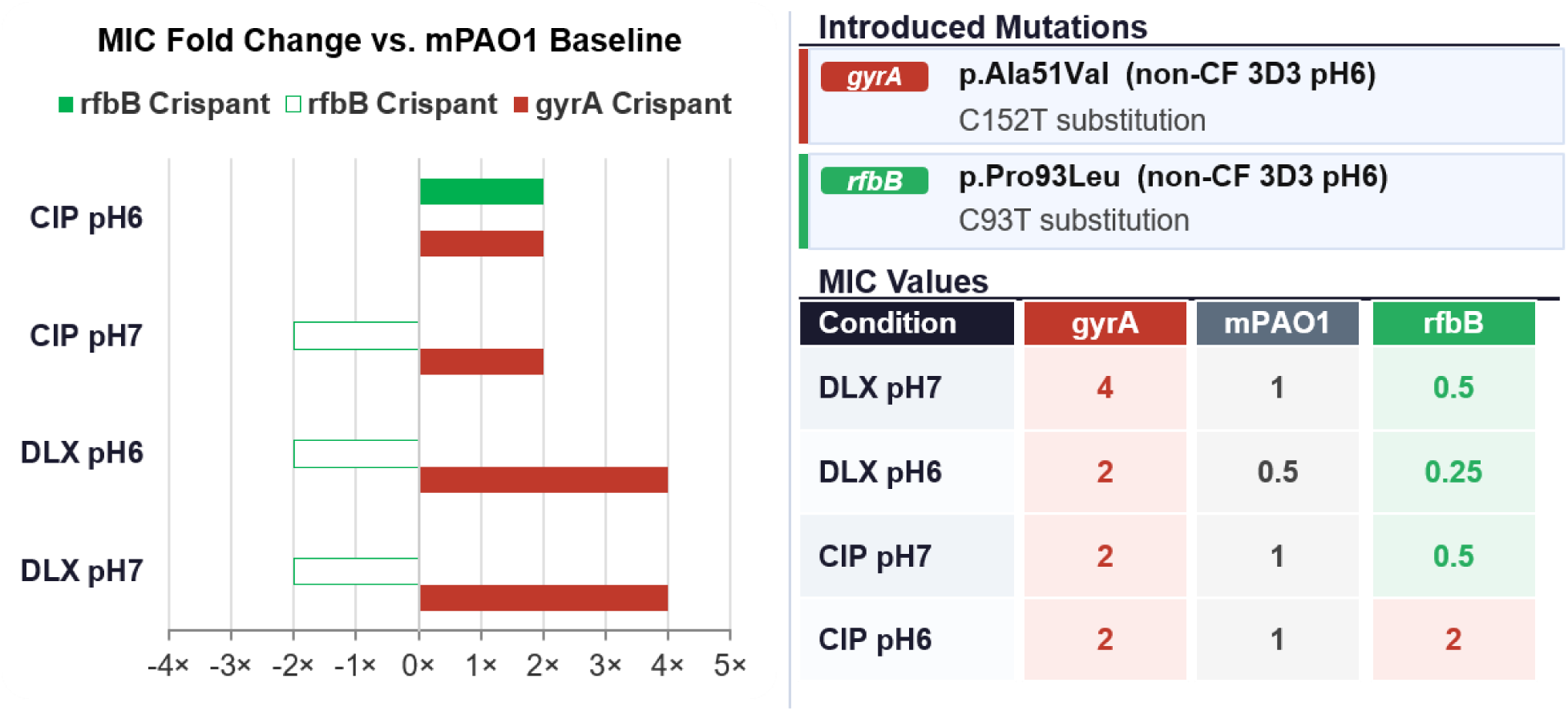
Diverging bar chart showing MIC fold change relative to the mPAO1 wild-type baseline for two CRISPR-edited strains across four antibiotic conditions. extending to the right indicate increased resistance, whereas hollow green bars extending to the left indicate increased susceptibility. The gyrA crispant (p.Ala51Val) showed a 2-to 4-fold increase in resistance to both DLX and CIP. The rfbB crispant (p.Pro93Leu) showed a twofold increase in susceptibility under most conditions but unexpectedly reverted to twofold resistance under CIP at pH 6.0. MIC values (mg/L) are provided in the accompanying table.

**Table 2:**
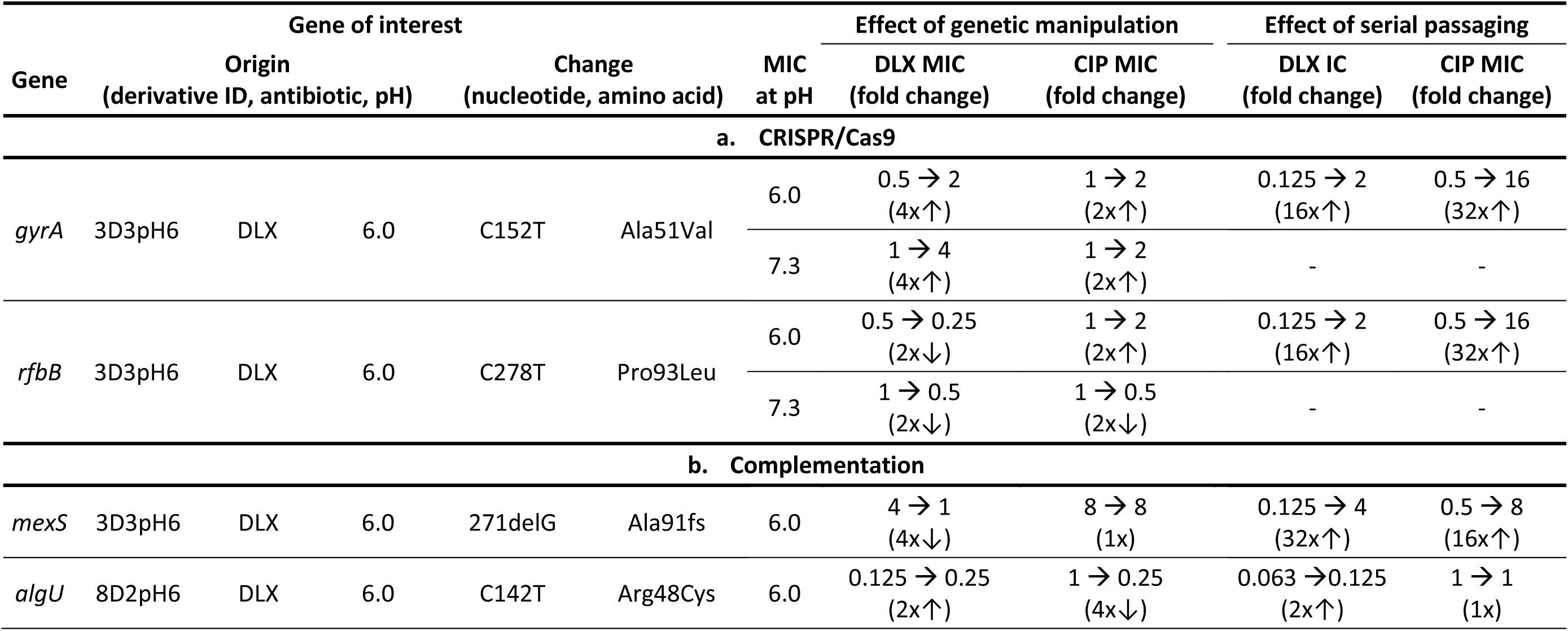
Genome engineering and functional validation of candidate genes for resistant derivatives derived under sub-inhibitory concentrations of DLX in pH 6.0 (a) CRISPR/Cas9: mutation introduced into mPAO1 to test whether it alone is sufficient to cause resistance. (b) Complementation: Introduced wild-type in the resistant derivative to test whether it is necessary for the resistance phenotype. Origin, the evolved derivative (isolate, selecting antibiotic, pH) from which the mutation was identified. Change, nucleotide substitution and resulting amino acid change. MIC at pH, pH at which the edited/complemented strain was tested. DLX MIC / CIP MIC (fold change), MIC before → after editing, with fold-change (↑/↓) in parentheses. Passaging effect (fold change), the DLX/CIP MIC fold-change originally observed in that lineage during serial passaging, shown for comparison; –, no matching passaging value (construct tested at a pH other than the lineage’s original passaging pH). DLX, delafloxacin; CIP, ciprofloxacin; MIC, Minimum inhibitory concentration (mg/L).

Despite repeated attempts, editing of *mexS* C270_271insC (p.Ala91fs) and *algU* G139C (p.Ala47Pro) could not be confirmed. These targets were therefore taken forward using complementation-based functional validation instead (**Figure 5a**).

**Figure 5:**
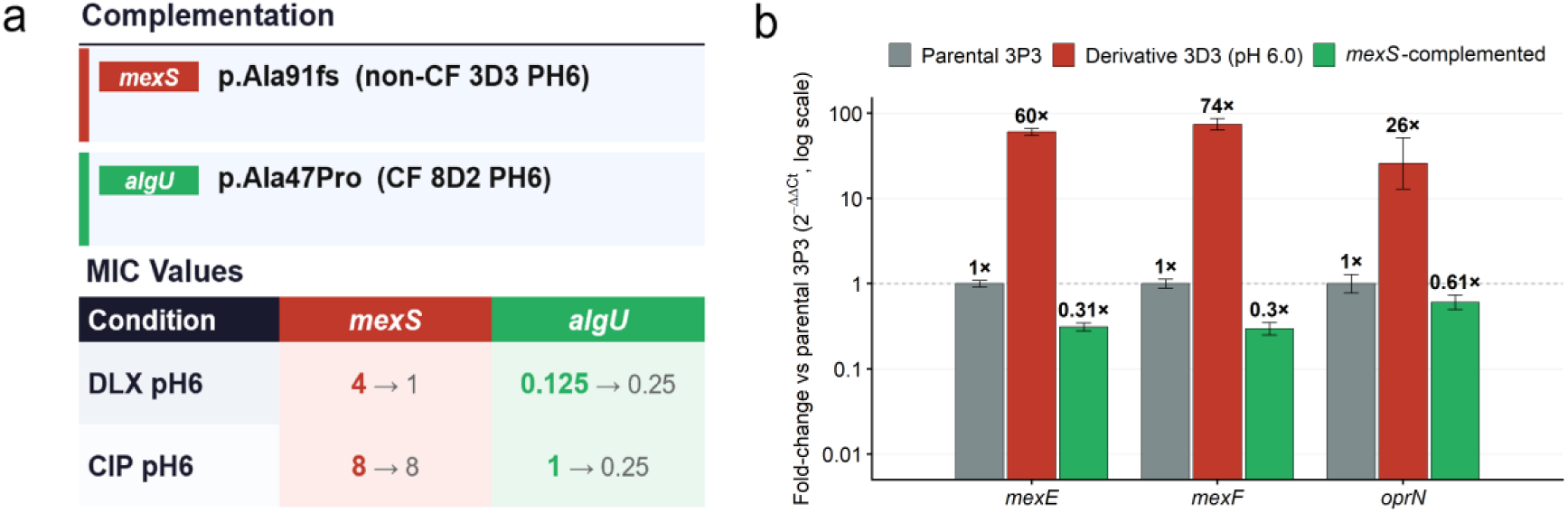
a) Complementation of the wild-type gene into the mutant background using pUCP24; mexS and algU complementation in P. aeruginosa strains 3D3 at pH 6.0 and 8D2 at pH 6.0. b) mexS complementation restored DLX susceptibility by repressing MexEF-OprN in the mexS-complemented 3D3 strain at pH 6.0. Fold-change is shown relative to the parental 3P3 strain, calculated as 2−ΔΔ*Ct* on a log scale. The reference gene was rpoD. Error bars represent ±1 SD propagated from technical triplicate Ct measurements through the ΔΔ*Ct*calculation. The dashed horizontal line indicates the parental baseline (1×).

qRT-PCR confirmed marked derepression of the MexEF-OprN operon in 3D3 pH 6.0 relative to the parental strain, with strong upregulation of *mexE* (60-fold), *mexF* (74-fold), and *oprN* (26-fold). Restoration of wild-type *mexS* in the 3D3 pH 6.0 background reduced DLX MIC fourfold, from 4 to 1 mg/L, without affecting CIP MIC **(Table 2b)** and introduction of functional *mexS* reversed this phenotype, reducing *mexE*, *mexF*, and *oprN* transcript levels to 0.31-, 0.30-, and 0.61-fold of parental, respectively (**Figure 5b**). Together, these data show that *mexS* loss-of-function drives DLX resistance through MexT-dependent MexEF-OprN overexpression, whereas CIP resistance is retained independently of *mexS* status.

Reintroduction of wild-type *algU* reduced CIP MIC fourfold, from 1 to 0.25 mg/L, and was accompanied by a smaller twofold increase in DLX MIC, from 0.125 to 0.25 mg/L (**Figure 5a**). Notably, this effect occurred in a strain without a prior resistance phenotype. This observation suggests a possible role for *algU* in modulating CIP susceptibility in this background, although causal inference will require reconstruction of the *algU* p.Ala47Pro allele in a clean mPAO1 background.

## Discussion

Our study demonstrates that delafloxacin exhibits a significantly higher genetic barrier to resistance evolution than ciprofloxacin in *Pseudomonas aeruginosa*, with particularly pronounced differences under acidic conditions that mimic the CF airway environment. These findings have direct implications for antibiotic stewardship in chronic pulmonary infections.

### DLX Maintains Potency and Resists Resistance Development Under Acidic Conditions

DLX retained significantly better activity than CIP under acidic conditions, and resistance evolved more slowly, consistent with earlier work in *E. coli* (*17*) but now shown here in *P. aeruginosa,* a clinically more challenging pathogen. In our experiment, resistance to DLX emerged in fewer lineages and with a smaller rise in MIC at pH 6.0 than at pH 7.3, and this pH effect was clear when the paired evolved lineages were split from the same clone into acidic and neutral conditions, were compared directly. Bösch et al. (17) reported a similar pH-dependent pattern in *E. coli*, and our data suggest that this phenomenon also applies to *P. aeruginosa*, despite its more complex resistance background, including multiple efflux systems and lower outer membrane permeability. Our data therefore suggest that acidic pH slows DLX resistance evolution in *P. aeruginosa* as well, even in a species with a more complex intrinsic resistance repertoire. Mechanistically, CIP resistance can often arise through single QRDR changes, whereas DLX appears to require a higher mutational burden, because escape from dual target inhibition is harder to achieve. That difference likely helps explain why DLX remained more robust against resistance development. (14, 17–20).

The clinical relevance is consistent with previous reports (10, 21) where a substantial fraction of CIP-resistant isolates remained susceptible to DLX, matching the partial separation we observed between CIP and DLX cross-resistance. This is also consistent with experimental and model-based data showing improved DLX performance in CF sputum and under CF-like acidic conditions, including superior time-kill activity, lower MICs relative to levofloxacin and CIP, and reduced activity of other antibiotics in acidic biofilm environments (11, 16). CF airway surface liquid is acidic in multiple independent studies, and biofilms can generate even lower local pH gradients (22–24). At acidic pH, DLX is expected to shift toward a more membrane-permeable form, which may promote intracellular accumulation and help explain the slower resistance evolution we observed. (25, 26).

### Novel gyrA Mutation Outside Canonical Hotspots Contributes to DLX Resistance

The *gyrA* p.Ala51Val substitution, although located outside the canonical Thr83/Asp87 QRDR hotspots (18, 19), was sufficient to confer DLX resistance in the mPAO1 background, increasing DLX MIC fourfold at both pH 7.3 and pH 6.0 and CIP MIC twofold. This supports a causal role for this noncanonical *gyrA* variant in fluoroquinolone resistance.

Atypical QRDR mutations are increasingly reported in CF-adapted *P. aeruginosa* under prolonged antibiotic exposure (27, 28). Our findings are consistent with the observation of Fujiwara et al., (29) who reported that *gyrA* mutations combined with *mexR*-deregulated MexAB-OprM efflux can produce very large fluoroquinolone resistance increases, reportedly up to 1024-fold. In our panel, *gyrA* p.Ala51Val also co-occurred with efflux-pathway activation, and these derivatives showed the highest MIC fold-rises, supporting a possible additive contribution of target-site and efflux-mediated resistance, although the magnitude of this interaction cannot be quantified from our data alone.

### MexEF-OprN Is Associated with pH-and Origin-Dependent Efflux Responses, Though Effects Are Trend-Level at this Sample Size

qRT-PCR profiling of *mexE*, *mexF*, and *oprN* in 24 resistant derivatives showed numerically higher induction under CIP selection in non-CF than CF isolates at pH 7.3, with a smaller gap at pH 6.0. None of the CF versus non-CF comparisons reached statistical significance, largely because the sample size per subgroup was only three isolates. Under DLX selection, all three genes tended to be more strongly induced at acidic pH irrespective of origin, whereas no significant origin effect was detected. Thus, pH appeared to be the stronger stratifying variable for DLX-driven MexEF-OprN induction, but the available data support this only as a trend, not a confirmed effect.

Restoration of wild-type *mexS* in the 3D3 pH 6.0 background reduced DLX MIC fourfold without affecting CIP MIC. This was accompanied by strong upregulation of *mexE* (60-fold), *mexF* (74-fold), and *oprN* (26-fold) in 3D3 pH 6.0, with reversion to subparental expression after *mexS* complementation, confirming that *mexS* loss-of-function drives DLX resistance through constitutive MexEF-OprN derepression (30–32).

This mechanism is compatible with the cross-resistance data, but it is not fully proven. Overall, DLX and CIP cross-resistance were not significantly different, but the asymmetry was clear at pH 6.0 and absent at pH 7.3. A reasonable explanation is that DLX more often selects efflux-based resistance, which tends to protect against both fluoroquinolones, whereas CIP preserved activity against CIP-selected populations could translate into a more durable therapeutic effect. At the same time, DLX is not resistance-neutral: exposure can select for CIP cross-resistance, mainly through MexEF-OprN overexpression, and therefore DLX should be used cautiously, particularly in chronic infections with prolonged or subinhibitory drug exposure.

## Materials and Methods

### Bacterial isolates and culture conditions

Six *P. aeruginosa* clinical isolates (3 cystic fibrosis [CF], 3 non-CF) from respiratory samples (bronchial/tracheal secretions, sputum, throat swab, bronchoalveolar lavage) were cultured in Mueller-Hinton broth (MHB, pH 7.3 or 6.0 adjusted with hydrochloric acid [HCl]) using Nunc™ CELLSTAR® 6-well plates (Greiner Bio-One GmbH, Frickenhausen, Germany) at 37°C with shaking (120–150 rpm) on a microplate shaker (Edmund Bühler GmbH, Germany).

DLX and CIP were purchased from Sigma-Aldrich Chemie GmbH, and stock solutions were made by dissolving powder stocks according to the manufacturer’s instructions in 0.2 µm filter-sterilized water and stored at −20°C before use. MHB was used at a standard pH of 7.3 and adjusted to an acidic pH of 6.0 by the dropwise addition of 1 M HCl (Sigma-Aldrich Chemie GmbH). pH measurement was performed with Seven Easy pH meter (Mettler-Toledo GmbH, Schwarzenbach, Switzerland).

### MIC determination / antimicrobial susceptibility testing

MICs for DLX and CIP were determined by the broth microdilution method ( antibiotic range 0.002–512 mg/L) in 96-well plates (Nunc CELLSTAR Greiner Bio-One GmbH, Frickenhausen, Germany) using a final volume of 100 µL per well and an inoculum of 5 × 10⁵ colony-forming unit (CFU)/mL with the broth pH adjusted to the pH that was used during the passages. MIC determination was done following the latest EUCAST recommendations and values were interpreted in accordance with the 2026 EUCAST criteria (version 16.1, 2026).

### Multi-step resistance selection

Three representative clones were selected from each of the six original *Pseudomonas aeruginosa* isolates by picking a single CFU and streaking it onto a fresh MH agar plate. From each plate, three distinct colonies were then chosen as parental clones for the subsequent multi-step resistance selection experiment. In total, 18 strains were passaged in each antibiotic condition (18 in CIP and 18 in DLX) at pH 6.0 and pH 7.3. Briefly, one single colony grown overnight at 37°C on an MH agar plate was picked and grown in 5 mL of MHB at 37°C without antibiotics (overnight culture). Fifty microliters of the overnight broth were transferred to fresh MHB (at the respective pH, 6.0 or 7.3) containing CIP or DLX at one-half of the clone’s MIC and incubated at 37°C with shaking (120-150 rpm). Twice daily, 50 µL of the previous culture was transferred to fresh MHB at the respective pH containing antibiotic at the same concentration, to correct for pH drift occurring during incubation. After 3 consecutive days at the same antibiotic concentration, MICs were reassessed by broth microdilution and the antibiotic concentration for the next passage was adjusted accordingly. Aliquots were sampled after each passage, washed with PBS to remove antibiotic, resuspended in MHB, and 25 µL was stored in 1 mL of 10% glycerol LB broth at −80°C for further molecular investigations. The same procedure was performed in parallel at both pH conditions without antibiotic pressure, serving as negative controls.

### Whole Genome sequencing & Mutation identification

Total DNA was extracted from the selected strains (*n* = 24 based on the condition and MICs) with the QiaAMP Mini Kit (QIAGEN, Hilden, Germany) following the manufacturer’s instructions. DNA concentrations were measured with a Nanodrop OneC spectrometer (Thermo Fisher Scientific, Massachusetts, USA).

WGS was performed at the Institute of Medical Microbiology, University of Zurich, under ISO 17025 accreditation. Library preparation used the QIASeq FX DNA Library Kit (Qiagen) and paired-end sequencing (2 × 150 bp) was performed on an Illumina NextSeq1000 platform (Illumina®, San Diego, CA, USA), ensuring a minimum average read depth of 30×. Quality control and assembly used the IMMense pipeline (https://gitlab.uzh.ch/appliedmicrobiologyresearch/immense). Variant calling used a pipeline involving read trimming with trimmomatic (33), mapping against an assembled parental strain with bwa (34), variant calling with pilon (35) and effects with SNPeff (36). Results were compared and visually checked against mapped reads in bam format in artemis (37).

### CRISPR/Cas9 genome editing

CRISPR/Cas9 genome editing was performed using the two plasmid (pCasPA/pACRISPR) method developed by Chen et al. (38) using the λ-Red recombination system; a comparable native CRISPR-Cas–mediated editing approach in clinical multidrug-resistant *P. aeruginosa* has been described previously (39).

sgRNA oligos were designed with CRISPOR and the Oligos were phosphorylated and annealed by mixing 37 µl H2O, 5 µl 10x T4 Polynucleotide Kinase Buffer (New England BioLabs, Ipswich, MA, USA), 5 µl fresh 10mM ATP, 1 µl 10U/µl T4 polynucleotide kinase (New England BioLabs(NEB), Ipswich, MA, USA), 1 µl of each oligo (100 µM). After one hour incubation at 37°C, 2.5 µl 1M NaCl were added, the mix heated to 95°C for 3 minutes and then cooled to 25°C by decreasing the temperature −0.1°C per second. Annealed oligos were diluted 1:20 and 1 µl mixed with 6 µl H2O, 1 µl 10x T4 DNA Ligase Buffer (NEB), 20nM pACRISPR plasmid, 0.5µl T4 DNA Ligase (400U/µl), 0.5 µl BsaI-HFv2 (NEB). Reaction was kept in a thermocycler for 25 cycles (37°C for 3 min, 16°C for 4 min), then 80°C for 15 min, and finally cooled to 10°C. The product was transformed into zymo mix and go (Takara, Mountain View, CA, USA) competent *E. coli* and selected on 50 mg/L carbenicillin plates. Insertion of the sgRNA sequence was verified by sequencing. Primer for the repair 200 bp arms were designed with the Takara In-Fusion Cloning Primer Design Tool to contain the desired restriction sites and containing the desired nucleotide change and a synonymous change of the PAM sequence. Repair arms were inserted in between the XhoI and XbaI restriction site of the pACRISPR plasmid following the Takara In-Fusion® Snap Assembly PCR Cloning systems protocol and insertion verified by PCR and sequencing. *P. aeruginosa* mPAO1 was made electrocompetent by diluting an overnight culture 1:100 in fresh LBB and incubating them at 37°C until OD600 of ∼0.5. Cells were chilled on ice for 30 minutes, then were harvested by centrifugation at 3300 rpm for 5 min and pellet washed twice with 20 ml of ice-cold 10% glycerol and then resuspended in 1 ml 10% glycerol. 100 µl electrocompetent cells were mixed with up to 5 µl of pCas plasmid and electroporated in a 2 mm cuvette (2.5kV, 200 Ω, 25 µF). Immediately after the pulse 1 ml LBB was added and the mix was incubated for 90 minutes at 37° C, 200 rpm followed by selection on 30 mg/L tetracycline plates.

Afterwards one colony harbouring the pCasPA plasmid was prepared as electrocompetent cells, same as above except for the addition of 2 mg/L total concentration L-arabinose at OD600 is 0.3. pACRISPR with the sgRNA sequence and repair arms were mixed and electroporation followed with the same conditions. 30 mg/L tetracycline and 100 mg/L carbenicillin were used for selection. Transformants were plasmid cured by growing them in the absence of antibiotics until evident growth and then plating a 1:10⁴ dilution on 5% sucrose LBA. The PCR amplified region of the CRISPR site of Plasmid cured bacteria was sequenced to confirm the CRISPR event.

### Complementation /cloning

To evaluate the impact of the mutations on DLX and CIP susceptibility, a series of cloning and expression experiments were performed. The candidate genes from parental isolates were PCR amplified with primers designed with the Takara In-Fusion Cloning Primer Design Tool to contain the desired restriction sites. The amplicons and the shuttle vector pUCP24 plasmid were digested with the corresponding restriction enzymes, purified, ligated with Quickligase (New England Biolabs, Ipswich, USA), and then transformed into competent *E. coli* (Top10 Mix & Go; Zymo Research, USA). For cloning *mexS* and *algU* into pUCP24 transformants were selected on 30 mg/L gentamycin LB agar plates treated with 2% X-gal and 0.1M isopropyl-β-D-thiogalactopyranosid (IPTG), the recombinant plasmids were extracted with the NucleoSpin Plasmid Kit (Macherey-Nagel, Düren, Germany), and the insertion was confirmed by sanger sequencing (Microsynth, AG, Balgach, Switzerland). Thereafter, they were transformed into the selected derivatives that were made electro competent and transformants were selected on 30 mg/L gentamycin LB agar plates. The DLX and CIP resistance of transformants was determined by broth microdilution. As a negative control, the shuttle vector pUCP24 without candidate genes was introduced to the strains to analyse the effect of the vector on the MICs of the antibiotics.

### Efflux pump gene expression analysis

Total RNA was extracted from exponentially growing cultures harvested at an optical density at 600 nm (OD₆₀₀) of 0.4–0.5 of all 24 resistant derivatives and the parental strain using RNAprotect Bacteria Reagent (QIAGEN, Hilden, Germany) for immediate transcriptional stabilisation, followed according to the RNAprotect Bacteria Reagent Handbook (QIAGEN). Expression of 14 efflux pump genes spanning the MexAB-OprM, MexCD-OprJ, and MexEF-OprN systems and their associated regulators was subsequently quantified by one-step qRT-PCR using 80 ng of total RNA per reaction with the RNA Power SYBR Green RNA-to-CT 1-Step Kit (Thermo Fisher Scientific) on the QuantStudio 5 Real-Time PCR System (Applied Biosystems). Relative expression was calculated by the ΔΔCT method using *rpoD* as the internal housekeeping reference gene according to (40). All values were normalised to the parental baseline and expressed as fold-change.

### Statistical analysis

Resistance was defined using species-specific breakpoints. EUCAST *P. aeruginosa* breakpoints were used for ciprofloxacin (CIP; resistant >0.5 mg/L), while FDA criteria were applied for delafloxacin (DLX; susceptible ≤0.5 mg/L, intermediate 1 mg/L, resistant ≥2 mg/L).

To account for the hierarchical design, both lineage-level and clone-paired analyses were performed. Fisher’s exact test treated all 36 lineages as independent, whereas exact McNemar’s test used 18 paired clone-level comparisons. MIC fold changes were analysed using Mann– Whitney U and paired Wilcoxon signed-rank tests and are presented as geometric means with 95% confidence intervals. CF and non-CF isolates were analysed as independent groups.

A GEE sensitivity analysis was performed for the DLX arm to account for clustering by parental strain. CIP was not modelled because all CIP-passaged lineages were resistant. Due to the limited number of parental strains and unstable variance estimates, GEE results were considered supportive only. Two-sided *p*<0.05 was considered statistically significant.

Geometric means, 95% confidence intervals and Mann-Whitney U comparisons were performed in GraphPad Prism version 10.2.2 (GraphPad Software, Boston, MA), which was also used to generate Figure 2. Fisher’s exact tests were performed using OpenEpi version 3.01 (41). McNemar’s exact tests were performed using GraphPad QuickCalcs Software (42). GEE models were fitted in R version 4.4.2 (R Core Team, 2024) using the geepack package version 1.3.13 (43) with Mancl-DeRouen small-sample corrected standard errors from the geesmv package version 1.3 (44).

## ACKNOWLEDGMENTS

This study was funded through a research grant from HOCH Health Ostschweiz, St. Gallen, Switzerland. Grant number 24/01

All other authors do not report any potential conflict of interest.

We thank the technicians of the IMM for their dedicated help and the University of Zurich for continuous support. This work made use of infrastructure services provided by the Science IT team of the University of Zurich (www.s3it.uzh.ch).

This study was presented at the 35th European Society of Clinical Microbiology & Infectious Diseases in Munich, Germany (Abstract No. 06387) and presenting at the Joint Annual Meeting of the Swiss Society for Infectious Diseases 2026 in Montreux, Switzerland (Abstract No.16209).

## DATA AVAILABILITY

The genomics data have been deposited in the European Nucleotide Archive (ENA) repository ENA Browser (https://www.ebi.ac.uk/ena/browser/text-search?query=PRJEB106985)

---

**Supplemental Table 1:** DLX and CIP MIC (mg/L) at day 0, 3, 6, and 9 of serial passaging for the 24 whole-genome-sequenced representative derivatives, with selecting antibiotic, pH, and CF/non-CF origin.

| Parental | Derivative | DLX MIC |  |  |  | CIP MIC |  |  |  | Culture condition |  |  |
| --- | --- | --- | --- | --- | --- | --- | --- | --- | --- | --- | --- | --- |
|  |  | Day 0 | Day 3 | Day 6 | Day 9 | Day 0 | Day 3 | Day 6 | Day 9 | Antibiotic | pH | CF/non-CF |
| 3P3 | 3D3_pH7.3 | 0.5 | 2 | 2 | 8 | 0.25 | 0.25 | 0.25 | 1 | DLX | 7.3 | non-CF |
| 5P2 | 5D2_pH7.3 | 0.25 | 1 | 1 | 16 | 0.25 | 0.25 | 2 | 1 | DLX | 7.3 | non-CF |
| 10P2 | 10D2_pH7.3 | 0.25 | 1 | 8 | 32 | 0.125 | 1 | 4 | 4 | DLX | 7.3 | non-CF |
| 3P1 | 3C1_pH7.3 | 0.5 | 2 | 8 | 64 | 0.25 | 4 | 8 | 256 | CIP | 7.3 | non-CF |
| 5P2 | 5C2_pH7.3 | 0.25 | 1 | 2 | 4 | 0.25 | 2 | 2 | 16 | CIP | 7.3 | non-CF |
| 10P3 | 10C3_pH7.3 | 0.25 | 0.25 | 1 | 2 | 0.125 | 0.25 | 2 | 8 | CIP | 7.3 | non-CF |
| 3P3 | 3D3_pH6.0 | 0.125 | 1 | 2 | 2 | 0.5 | 8 | 8 | 16 | DLX | 6 | non-CF |
| 5P2 | 5D2_pH6.0 | 0.063 | 0.031 | 0.5 | 0.25 | 0.25 | 0.5 | 4 | 8 | DLX | 6 | non-CF |
| 10P2 | 10D2_pH6.0 | 0.063 | 0.25 | 0.5 | 1 | 0.25 | 1 | 2 | 8 | DLX | 6 | non-CF |
| 3P2 | 3C2_pH6.0 | 0.125 | 1 | 2 | 32 | 0.5 | 4 | 64 | 512 | CIP | 6 | non-CF |
| 5P3 | 5C3_pH6.0 | 0.063 | 0.25 | 4 | 4 | 0.25 | 8 | 16 | 256 | CIP | 6 | non-CF |
| 10P3 | 10C3_pH6.0 | 0.063 | 0.125 | 2 | 4 | 0.25 | 2 | 8 | 32 | CIP | 6 | non-CF |
| 8P2 | 8D2_pH7.3 | 0.5 | 2 | 2 | 16 | 0.5 | 2 | 2 | 16 | DLX | 7.3 | CF |
| 9P1 | 9D1_pH7.3 | 0.5 | 2 | 16 | 32 | 0.125 | 1 | 16 | 16 | DLX | 7.3 | CF |
| 11P1 | 11D1_pH7.3 | 0.125 | 1 | 2 | 16 | 0.25 | 2 | 4 | 16 | DLX | 7.3 | CF |
| 8P3 | 8C3_pH7.3 | 0.5 | 4 | 16 | 64 | 0.5 | 8 | 32 | 256 | CIP | 7.3 | CF |
| 9P1 | 9C1_pH7.3 | 0.5 | 4 | 8 | 32 | 0.125 | 2 | 8 | 32 | CIP | 7.3 | CF |
| 11P3 | 11C3_pH7.3 | 0.125 | 1 | 2 | 16 | 0.25 | 2 | 8 | 64 | CIP | 7.3 | CF |
| 8P2 | 8D2_pH6.0 | 0.063 | 0.031 | 0.125 | 0.063 | 1 | 0.5 | 1 | 1 | DLX | 6 | CF |
| 9P3 | 9D3_pH6.0 | 0.063 | 1 | 4 | 4 | 1 | 0.5 | 16 | 16 | DLX | 6 | CF |
| 11P2 | 11D2_pH6.0 | 0.016 | 0.031 | 0.25 | 0.25 | 1 | 2 | 16 | 16 | DLX | 6 | CF |
| 8P3 | 8C3_pH6.0 | 0.063 | 0.063 | 0.063 | 0.25 | 1 | 0.5 | 1 | 4 | CIP | 6 | CF |
| 9P1 | 9C1_pH6.0 | 0.063 | 2 | 4 | 16 | 1 | 2 | 16 | 64 | CIP | 6 | CF |
| 11P3 | 11C3_pH6.0 | 0.016 | 0.125 | 0.5 | 8 | 1 | 8 | 128 | 512 | CIP | 6 | CF |

**Supplemental Table 2:** Efflux pump gene expression (RT-qPCR fold-change vs. parental strain) and corresponding DLX/CIP MIC (mg/L) in day 9 derivatives, sorted by selecting antibiotic and pH. -, alue not determined.

| Derivative | Antibiotic | pH | *mexA* | *mexB* | *oprM* | *nalD* | *mexR* | *mexC* | *mexD* | *nalC* | *oprJ* | *nfxB* | *mexT* | *oprN* | *mexE* | *mexF* | DLX MIC | CIP MIC |
| --- | --- | --- | --- | --- | --- | --- | --- | --- | --- | --- | --- | --- | --- | --- | --- | --- | --- | --- |
| 3C2 pH6 | CIP | 6.0 | 0.821 | 0.458 | 0.970 | 0.645 | 0.807 | - | 1.952 | 1.806 | 1.852 | 1.696 | 1.611 | 14.199 | 55.772 | 133.331 | 32 | 512 |
| 5C3 pH6 | CIP | 6.0 | 0.609 | 0.646 | 0.879 | 0.592 | 0.662 | 0.483 | 0.637 | 0.629 | 0.507 | 0.479 | 0.681 | 6.671 | 33.324 | 19.088 | 4 | 256 |
| 10C3 pH6 | CIP | 6.0 | 1.115 | 0.696 | 0.843 | 0.518 | 0.976 | 1.610 | 1.297 | 0.920 | 0.955 | 1.029 | 1.298 | 45.805 | 133.555 | 278.307 | 4 | 32 |
| 8C3 pH6 | CIP | 6.0 | 0.241 | 0.271 | 0.351 | 0.173 | 0.417 | 0.579 | 1.012 | - | 0.866 | 0.768 | 0.474 | 2.126 | 13.526 | 21.517 | 0.25 | 4 |
| 9C1 pH6 | CIP | 6.0 | 0.504 | 0.843 | 1.246 | 1.246 | 1.349 | 0.627 | 0.918 | 2.034 | 1.027 | 1.099 | 2.081 | 8.942 | 5.585 | 5.813 | 16 | 64 |
| 11C3 pH6 | CIP | 6.0 | 1.026 | 1.108 | 0.507 | 0.750 | 0.945 | 2.361 | 6.860 | 0.870 | 4.004 | 2.436 | 0.679 | 1.089 | 1.388 | 1.226 | 8 | 512 |
| 3D3 pH6 | DLX | 6.0 | 0.506 | 0.714 | 1.215 | 0.509 | 0.879 | - | 2.180 | 0.673 | 0.836 | 1.590 | 0.885 | 58.590 | 77.008 | 9.425 | 2 | 16 |
| 5D2 pH6 | DLX | 6.0 | 0.961 | 0.804 | 0.881 | 0.676 | 0.971 | 1.580 | 1.222 | 0.853 | 2.018 | 1.147 | 2.040 | 6.582 | 31.988 | 25.442 | 0.25 | 8 |
| 10D2 pH6 | DLX | 6.0 | 0.836 | 0.794 | 1.028 | 0.576 | 1.112 | 0.938 | 1.036 | 0.584 | 0.561 | 0.656 | 1.063 | 24.232 | 80.596 | 114.163 | 1 | 8 |
| 8D2 pH6 | DLX | 6.0 | 0.999 | 0.870 | 1.160 | 0.582 | 0.909 | 0.629 | 0.690 | - | 0.615 | 0.751 | 0.714 | 0.609 | 0.502 | 0.640 | 0.063 | 1 |
| 9D3 pH6 | DLX | 6.0 | 4.956 | 2.041 | 2.633 | 0.898 | 2.882 | 1.303 | 1.040 | 2.059 | 0.761 | 0.854 | 0.679 | 17.031 | 58.190 | 106.866 | 4 | 16 |
| 11D2 pH6 | DLX | 6.0 | 1.611 | 0.541 | 1.306 | 1.402 | 1.014 | 3.415 | 10.705 | 1.657 | 8.122 | 5.089 | 1.653 | 2.274 | 2.237 | 2.219 | 0.25 | 16 |
| 3C1 pH7 | CIP | 7.3 | 0.912 | 0.595 | 0.859 | 0.571 | 0.900 | - | 0.822 | 1.120 | 0.419 | 1.406 | 2.578 | 14.222 | 196.680 | 335.907 | 64 | 256 |
| 5C2 pH7 | CIP | 7.3 | 1.123 | 0.900 | 1.151 | 0.548 | 0.569 | 0.916 | 1.061 | 1.056 | 1.142 | 1.437 | 1.405 | 39.073 | 51.168 | 46.903 | 4 | 16 |
| 10C3 pH7 | CIP | 7.3 | 1.112 | 0.554 | 0.566 | 0.505 | 0.777 | 1.025 | 1.045 | 1.259 | 0.611 | 0.783 | 1.016 | 28.802 | 60.019 | 122.682 | 2 | 8 |
| 8C3 pH7 | CIP | 7.3 | 0.810 | 0.575 | 0.519 | 0.754 | 0.957 | 8.889 | 12.595 | - | 8.344 | 4.728 | 0.783 | 0.749 | 0.988 | 1.158 | 64 | 256 |
| 9C1 pH7 | CIP | 7.3 | 0.419 | 0.560 | 0.863 | 1.420 | 1.103 | 0.447 | 1.576 | 1.610 | 0.655 | 1.265 | 2.338 | 4.698 | 3.585 | 4.134 | 32 | 32 |
| 11C3 pH7 | CIP | 7.3 | 0.959 | 0.937 | 0.568 | 0.950 | 0.676 | 1.304 | 1.886 | 1.044 | 1.776 | 1.361 | 1.200 | 0.784 | 1.926 | 1.508 | 16 | 64 |
| 3D3 pH7 | DLX | 7.3 | 1.939 | 3.236 | 2.988 | 0.276 | 1.274 | - | 0.961 | 0.773 | 1.080 | 1.032 | 0.627 | 0.645 | 0.515 | 0.593 | 8 | 1 |
| 5D2 pH7 | DLX | 7.3 | 2.758 | 4.251 | 2.681 | 1.189 | 2.097 | 2.038 | 1.504 | 1.573 | 1.429 | 1.488 | 1.996 | 1.349 | 2.413 | 2.060 | 16 | 1 |
| 10D2 pH7 | DLX | 7.3 | 2.536 | 2.019 | 2.798 | 0.797 | 1.268 | 0.894 | 1.113 | 0.799 | 0.722 | 0.830 | 0.770 | 1.151 | 2.930 | 2.657 | 32 | 4 |
| 8D2 pH7 | DLX | 7.3 | 0.134 | 0.463 | 0.661 | 0.997 | 0.589 | 2.185 | 3.800 | - | 4.873 | 3.304 | 0.395 | 0.293 | 0.377 | 0.496 | 16 | 16 |
| 9D1 pH7 | DLX | 7.3 | 4.024 | 2.799 | 5.128 | 1.084 | 2.180 | 0.989 | 2.783 | 3.143 | 0.622 | 0.944 | 1.404 | 1.004 | 1.518 | 3.227 | 32 | 16 |
| 11D1 pH7 | DLX | 7.3 | 15.969 | 68.676 | 20.794 | 0.570 | 15.483 | 2.688 | 3.385 | 7.553 | 8.846 | 1.445 | 5.026 | 2.578 | 6.054 | 6.289 | 16 | 16 |

**Supplemental Table 3:**
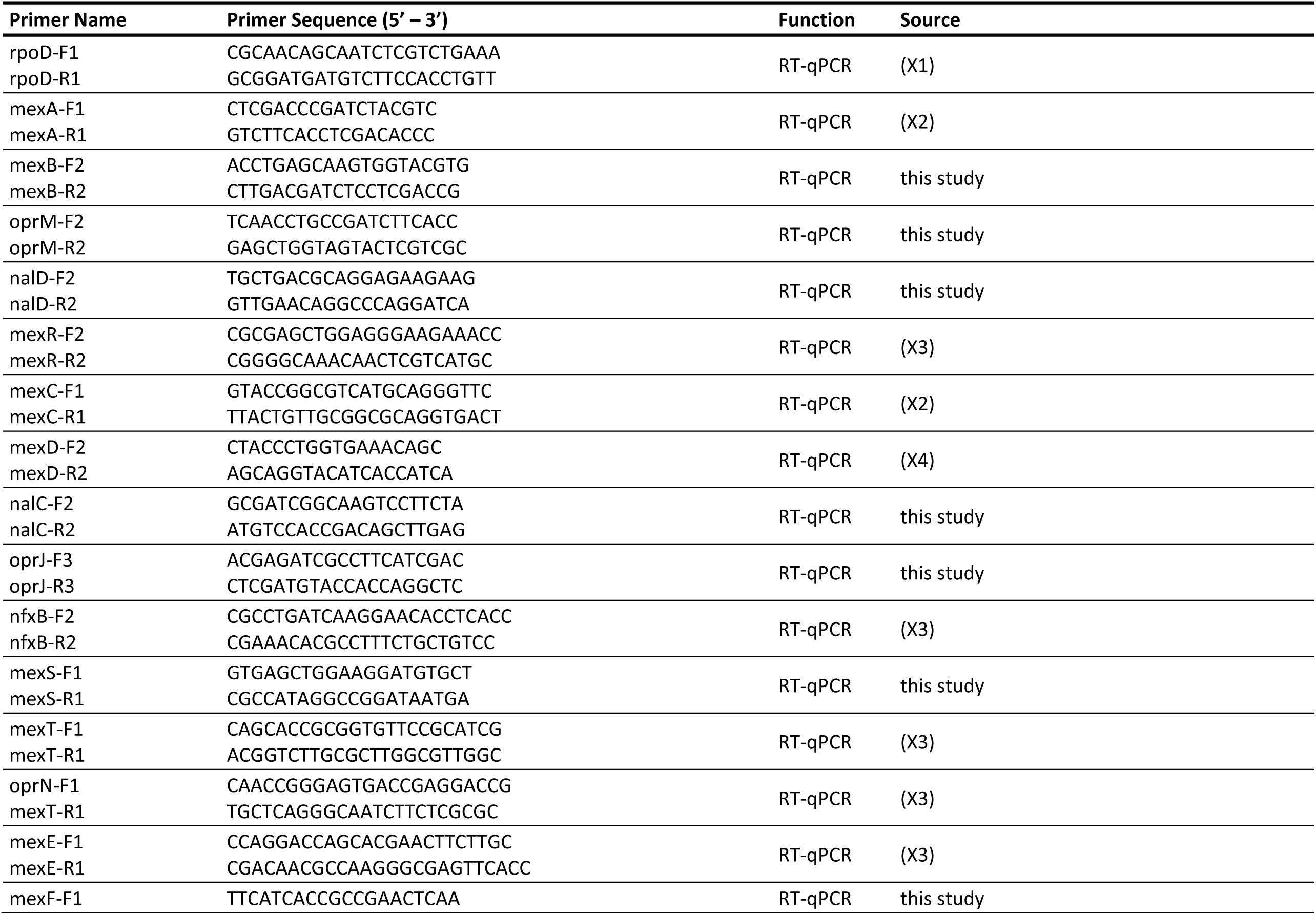
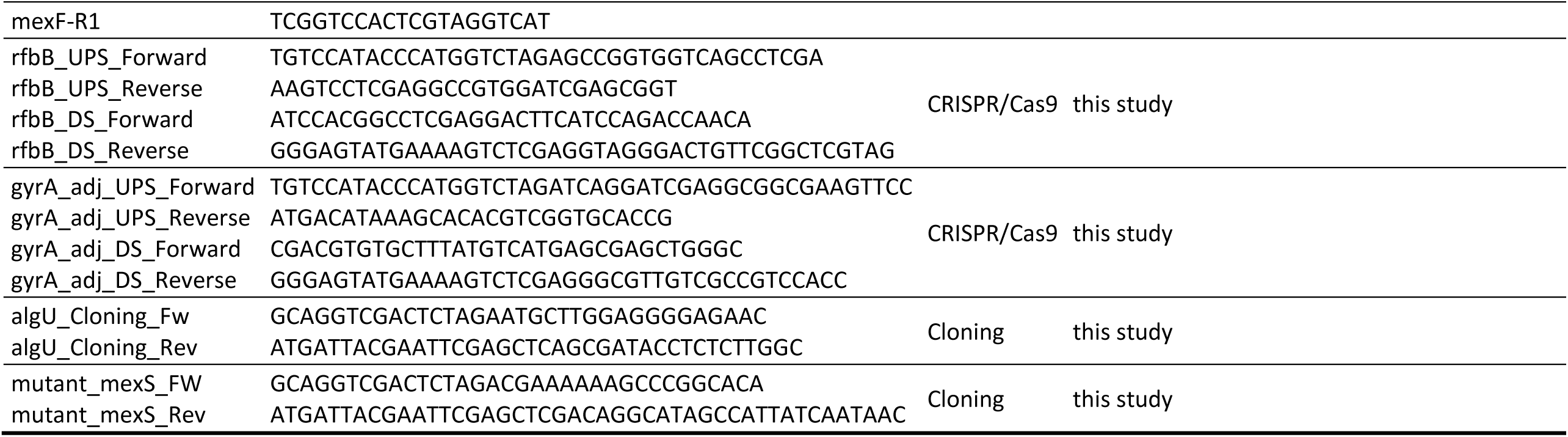
Primers used for RT-qPCR quantification of efflux pump gene expression (Supplemental Table 2), CRISPR/Cas9 genome engineering and complementation cloning (Table 2).

## Notes

### Competing Interest Statement

The authors have declared no competing interest.

